# Identification and Analysis of SARS-CoV-2 Mutation and Subtype using 2x tiled Primer Set with Oxford Nanopore Technologies Sequencing for Enhanced Variant Detection and Surveillance in Seoul, Korea

**DOI:** 10.1101/2023.12.03.569831

**Authors:** Giyoun Han, Sojung Lee, YaeEun Kwon, Jaemyun Lyu, Hyunsoo Kim, Kang-Jun Yoon, Minlee Kim

**Affiliations:** R&D Center. Genolution, Inc. 63, Magokjungang 8-ro 3-gil, Gangseo-gu, Seoul, Republic of Korea; Saint Peter’s Hospital. 2649, Nambusunhwan-ro, Gangnam-gu, Seoul, Republic of Korea

**Author notes:** **[Corresponding author]** Minlee Kim.

**Keywords:** Amplicon, Mutations, Next-generation sequencing, SARS-CoV-2, Long read sequencing

## Abstract

Severe acute respiratory syndrome coronavirus 2 (SARS-CoV-2) is a respiratory virus that contains RNA as its genetic material and has caused a global pandemic since its outbreak in 2020. This virus has many mutations, some of which can reduce the effectiveness of existing vaccines. Therefore, next-generation sequencing (NGS) is necessary to accurately identify new mutations. Current NGS analysis of SARS-CoV-2 uses the amplicon analysis method through a multiplex polymerase chain reaction. This study collected and validated RNA samples from patients who tested positive for SARS-CoV-2 from April to July 2022, and selected 613 samples for sequencing. The findings demonstrate the importance of long-read-based NGS analysis and 2x tiled primer set for identifying full SARS-CoV-2 genome sequence with new mutations and understanding the correlation between viral genotypes and patient characteristics for the effective management of SARS-CoV-2.

## 1. Introduction

Coronavirus disease, caused by severe acute respiratory syndrome coronavirus 2 (SARS-CoV-2), was declared a global pandemic by the World Health Organization (WHO) in March 2020 and had caused more than 460 million confirmed cases and more than 6.9 million deaths by March 2023 [1]. SARS-CoV-2 uses RNA as its genetic material. RNA viruses are easily mutated during gene replication, and many mutations occur within a short period [2, 3]. Usually, these mutations are ineffective; however, in rare cases, they create highly contagious and lethal variants, causing serious medical problems. In addition, spikes and surface proteins play important roles in infection and serve as target sites for vaccines [4, 5]. Mutations in these proteins reduce the efficacy of existing vaccines [6-8].

The WHO and U.S. Centers for Disease Control and Prevention closely monitor the occurrence of these mutant viruses and designate variants with high lethality and transmissibility. To date, variants ranging from alpha to omicron have been classified as variants of concern and variants of interest [9].

Most current quantitative polymerase chain reaction (qPCR) tests target the RdRp, N, and E genes [10, 11]. These genic regions are generally less mutated and can be used to confirm SARS-CoV-2, even if it has some mutations. However, S genes that generate various mutations are difficult to utilize as qPCR targets, and if the mutation site overlaps with the PCR primer site, false negatives may occur, hindering detection of new mutations.

Therefore, it is impossible to accurately identify new mutations without next-generation sequencing (NGS). Current SARS-CoV-2 NGS analysis uses an amplicon-based enrichment method through multiplex PCR [12, 13]. Typically, the primer set used for Illumina analysis (COVIDSeq Test) generates 98 amplicons, and the size of each amplicon is about 400 bp, whereas the primer set used for Oxford Nanopore Technologies (ONT) analysis (Midnight Amplicon panel) generates only 29 amplicons with 1,200 bp [14, 15]. Amplicons cannot be generated if the primer site is mutated, and this is less likely to occur in ONT analysis because of the larger amplicon size and fewer amplicon counts than in Illumina analysis. While the base quality of ONT is inferior to that of Illumina, multiplex PCR amplicon-based analysis can generate sufficient reads to compensate for some low base quality in both analysis methods [16, 17]. In addition, ONT analysis takes 8–24 h, whereas Illumina sequencing requires approximately 40 h or more [18].

We implemented 2x tiled primer sets for better coverage of SARS-CoV-2 genome sequences and adopted ONT to sequence SARS-CoV-2 genotypes using specimens from SARS-CoV-2 positive volunteers, living in the Seoul metropolitan area, South Korea. We also examined the association between clinical symptoms and SARS-CoV-2 subtypes, especially BA.2 and BA.5.

## 2. Materials & Methods

### 2.1 Ethics

All human samples in this study were obtained from voluntarily participating patients with written informed consent. The research plan and processes were reviewed and approved by the Institutional Review Board of Wiltse Memorial Hospital (2022-W07).

### 2.2 Sample collection and validation

Specimens were collected from patients who tested positive for SARS-CoV-2 between April and July 2022 using nasopharyngeal swabs at Saint Peter’s Hospital, Seoul, Republic of Korea. The viral RNA was extracted using Nextractor and VN kit (Genolution, Seoul, Republic of Korea) and validated with qRT-PCR using STANDARD M nCoV Real-Time Detection Kit (SD BioSensor, Suwon, Republic of Korea). Samples with a Ct value of 30 or less were selected for sequencing, resulting in a total of 613 samples.

### 2.3 cDNA synthesis and multiplex PCR for SARS-CoV-2-specific amplification

cDNA was synthesized from extracted viral RNA using the LunaScript® RT SuperMix Kit (NEB, Ipswich, MA) according to the manufacturer’s protocol. Synthesized cDNA was used as a template for SARS-CoV-2-specific multiplex PCR using xGen™ SARS-CoV-2 Midnight Amplicon Panel (IDT, USA) containing two primer pools designed for the emerging SARS-CoV-2 variants. Pools A and B included odd and even region primers, respectively, and were used to generate overlapping amplicons approximately 1,200 bp in length. PCR using primer pools A and B was performed on separate plates to avoid overlapping amplicons. PCR was conducted using a Rapid Barcoding Kit 96 (ONT, Oxford, UK) with Q5® Hot Start High-Fidelity 2X Master Mix (NEB, Ipswich, MA) according to the manufacturer’s protocol designed for SARS-CoV-2.

### 2.4 Designing 2x tiled Midnight primer sets for amplification failure prevention

An additionally designed primer C/D set was developed to improve the coverage of ONT sequencing using the Midnight primer (primer A/B) set. The primer C/D set comprised 15 and 14 pairs of primers for C and D, respectively, and produced an approximately 1,200 bp amplicon. It avoids overlapping regions with the existing sets and considers factors, such as primer redundancy, self-dimer formation, non-target (human genomic DNA) ratio, and amplicon size during primer design. We optimized the efficiency of the primer set by redesigning and adding primers to the set. Finally, 2x tiled primer set (Midnight primer + primer C/D) was produced to process complementarily with mitigating unexpected drop-out region due to primer site mutation based mis-amplification.

### 2.5 Library preparation and sequencing

The barcoding process involved pooling amplicons A and B from each sample. The pooled amplicons were then mixed with one–1-96 rapid barcodes from the Rapid Barcoding Kit 96 (ONT, Oxford, UK). The mixture was incubated at 30°C for 2 min and then at 80°C for 2 min. The barcoded samples were pooled and purified according to the manufacturer’s protocol. The library was prepared by combining purified library with Rapid Adapter F from ONT native and incubating at room temperature (24 °C) for 5 min. The library was loaded onto a MinION sequencer using an R 9.4.1 flow cell (ONT, Oxford, UK).

### 2.6 SARS-CoV-2 genotyping and viral subtype classification

The sequenced reads were based on guppy2 and processed using the poreCov (v1.5.1) [19] and epi2me-labs/wf-artic (v0.3.11) tools. Using the two analysis tools, read quality, filtered FASTQ and BAM, consensus sequence, lineage (pangolin, scorpio), clade (Nextclade), and genome coverage were analyzed. The more detailed and diverse poreCov results were used as the main results, and the epi2me-labs/wf-artic analysis results were used for crosschecking.

The results from poreCov and epi2me-labs/wf-artic tools were compared. Most of the mutations were detected by both tools, but some position-specific mutations were detected disconcordantly. For exclusive mutations, we inspected them with Integrative Genomics Viewer [20] to determine whether they were true or false.

To identify the subtype of complex recombinants, Lineage deComposition for SARS-CoV-2 was also performed within a poreCov pipeline [21].

### 2.7 Statistical Analysis

Patient characteristics, including age, sex, and symptoms, were collected and combined with SARS-CoV-2 genotyping and subtype classification results. For statistical analysis, the features were analyzed by t-tests for age-related comparisons with inhouse python code.

## 3. Results

### 3.1 Improved detection of variants with 2x tiled primer sets

Compared to conventional primer set (primer set A/B of Midnight primer), 2x tiled primer set (primer set A+C / B+D) yielded better results for a range of mutations, including Omicron, Mu, and earlier strains such as 19A and 20A (Figure 1). These findings highlight the superiority of our approach over the existing methods and provide important insights into the detection of viral mutations. The 2x tiled primer set may enhance the detection and monitoring of the spread of SARS-CoV-2 and its variants.

**Figure 1.**
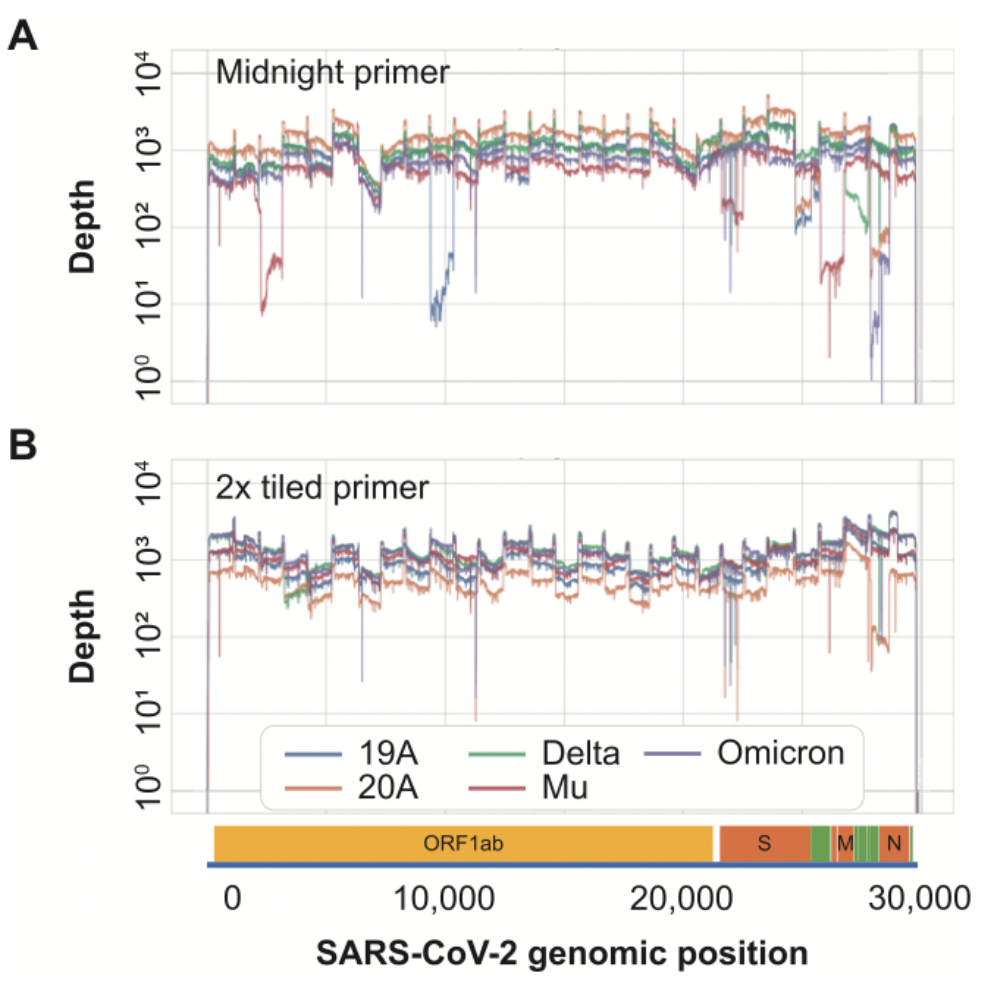
Covered depth by SARS-CoV-2 subtypes. Covered depth amplified with single primer set (A) and 2x tiled primer set (B) according to SARS-CoV-2 genome and genes. SARS-CoV-2 subtypes are colored as follows: 19A (blue), 20A (orange), Delta (green), Mu (red), and Omicron (purple). SARS-CoV-2, severe acute respiratory syndrome coronavirus 2, ORF, open reading frame, S, spike glycoprotein, M, membrane protein, N, nucleocapsid protein.

We also identified large deletions in SARS-CoV-2 by modifying the composition of the primer set. By designing a set capable of amplifying approximately 3,600 bp, in addition to the standard 1,200 bp (Supplementary Figure 1), we were able to successfully amplify previously failed regions.

### 3.2 Lineage and single-nucleotide variation (SNV) Analysis of Omicron Mutations

In this study, all analyzed SARS-CoV-2 sequences were reported to GISAID and all samples were classified as the omicron subtype: the BA.2 Omicron subtype (n=506) and BA.5 Omicron subtype (n=87). Additionally, there were 6 cases of the BA.1-like subtype, 2 cases of the BA.4-like subtype, and 12 cases of other variant subtypes. BA.2 was detected throughout the period of sample collection, while BA.5 was first found in May and had more confirmed cases than the BA.2 subtype in July. This pattern was also similar to that of the SARS-CoV-2 lineage type in Korea registered with the GISAID during the same period (April to July 2022) (Supplementary Figure 2).

We covered the full SARS-CoV-2 genome and identified a set of SNVs unique to the BA.2 and BA.5 lineages. The locations of these BA.2 and BA.5 specific SNVs were consistent with publicly available data, including the Cov lineage and NextClade, implying that although ONT sequencing has low base quality, its results can be used to detect mutations and classify viral subtypes (Figure 2).

**Figure 2.**
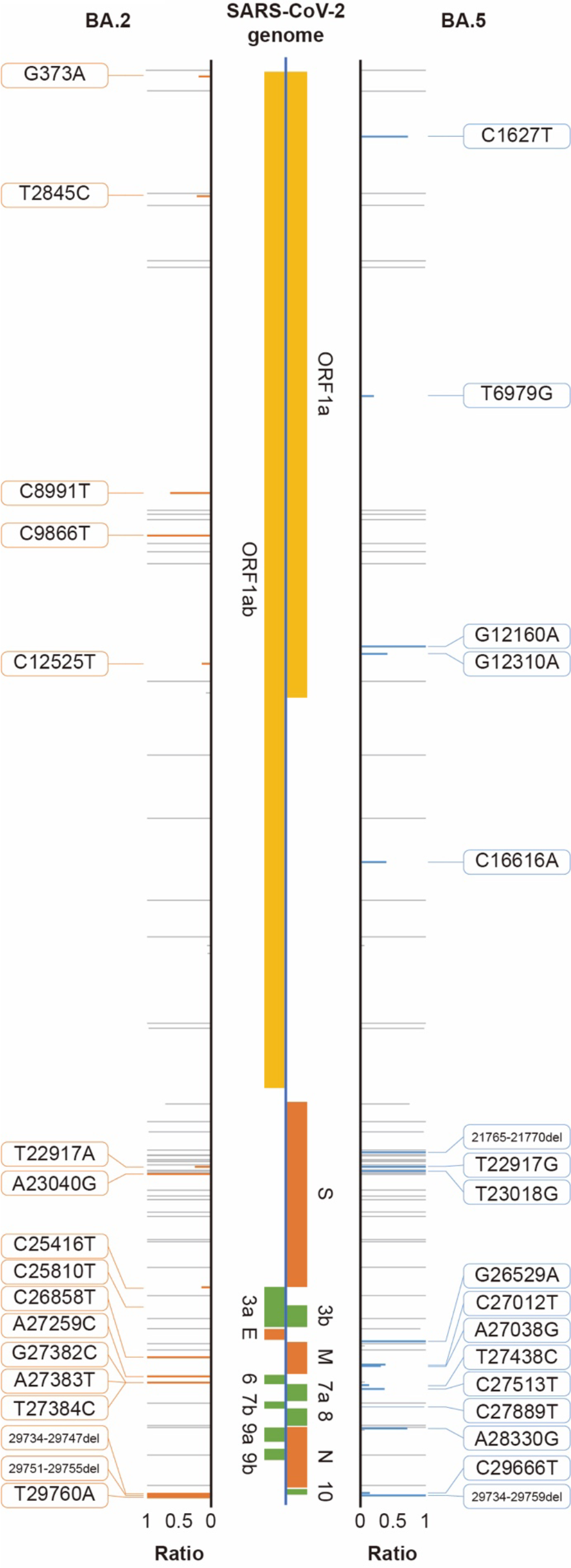
Subtype-specific mutations and ratios between BA.2 and BA.5 subtypes. Identified mutations for each subtype and mutation frequency ratio displayed with orange (BA.2) and blue (BA.5) and grey for common mutation.

During the analysis, the subtypes of some samples were not properly identified. In one sample identified as a recombinant sample, although it had an Omicron mutation, a clear constellation and sublineage could not be identified. Lineage deComposition for SARS-CoV-2 analysis revealed that the sample was a mutant in which BA.1 and BA.2 were recombined (Supplementary Figure 3) and later named as XE variant (2022-01-19, recombinant lineage of BA.1 and BA.2) [22]. Furthermore, no amplicons were generated in six samples, even though 2x tiled primer sets were used. In these cases, we found that 2–3 amplicons were not generated among the 29 amplicon pools. We modified the primer composition for detecting the large deletion at that location and sequenced again by separately applying a primer set covering the unamplified region. Separate sequencing results were concatenated and reanalyzed (Supplementary Figure 4). We sequenced and identified 936 bp, 929 bp, 851 bp, and 425 bp deletion in the SARS-CoV-2 genome. There were three large deletions, which included ORF7a, ORF7b, and ORF8 genic regions, and 425 bp deletion overlapped with ORF8. These SARS-CoV2 variants were also reported in GISAID (EPI_ISL_14435402, EPI_ISL_14467955, EPI_ISL_15269593, EPI_ISL_17463497).

### 3.3 SARS-CoV-2 subtype information and clinical characteristics of patients

Table 1 summarizes the viral variants and clinical characteristics of 613 individuals infected with SARS-CoV-2. The majority of cases had BA.2 subtype (n=506, 82.6%), followed by BA.5 (n=87, 14.2%). The mean age of the participants was 40.3 years (standard deviation=20.4), with slightly more females (n=321) than males (n=292); however, there was no statistically significant difference in the severity of infection or symptom manifestation between the two sexes. The most commonly reported symptoms were sore throat (n=362; 59.1%), persistent cough/sneezing (n=262; 42.7%), and sputum production (n=171; 27.9%). Fever (n=164, 26.8%), headache, muscle aches, chills, and runny nose were also reported by a significant proportion of the sample. Asymptomatic cases were relatively rare (4.4 %; n = 27). Due to small sample counts, data for some variants, such as BA.1 and BA.4, were excluded from the subsequent analysis.

**Table 1.** Characteristics and counts of patients by subtype, sex, age, and symptom.

| Characteristic | Total lineage | BA.1 | BA.2 | BA.4 | BA.5 | etc. |
| --- | --- | --- | --- | --- | --- | --- |
| Total no. | 613 | 6 | 506 | 2 | 87 | 12 |
| Sex |  |  |  |  |  |  |
| M | 292 | 4 | 242 | 1 | 39 | 6 |
| F | 321 | 2 | 264 | 1 | 48 | 6 |
| Mean age, y (SD) | 40.3 (20.4) | 55.2 (15.5) | 41.3 (20.2) | 17.5 (14.8) | 35.0 (20.0) | 31.7 (20.4) |
| Any Symptom |  |  |  |  |  |  |
| Asymptomatic | 27 | 0 | 23 | 0 | 2 | 2 |
| Sore throat | 362 | 2 | 305 | 2 | 48 | 5 |
| Persistent cough/sneezing | 262 | 4 | 219 | 1 | 33 | 5 |
| Sputum | 171 | 1 | 136 | 1 | 30 | 3 |
| Fever | 164 | 0 | 136 | 0 | 26 | 2 |
| Headache | 110 | 0 | 85 | 1 | 22 | 2 |
| Muscle aches | 99 | 1 | 80 | 0 | 17 | 1 |
| Chills | 80 | 1 | 61 | 0 | 15 | 3 |
| Runny nose | 73 | 1 | 63 | 0 | 9 | 0 |
| Hoarseness | 9 | 0 | 9 | 0 | 0 | 0 |
| Sickness | 8 | 0 | 7 | 0 | 1 | 0 |
| Low fever | 8 | 0 | 7 | 0 | 1 | 0 |
| Diarrhea | 7 | 0 | 7 | 0 | 0 | 0 |
| Dizziness | 6 | 0 | 6 | 0 | 0 | 0 |
| Chest pain | 4 | 0 | 4 | 0 | 0 | 0 |
| Nausea | 3 | 0 | 3 | 0 | 0 | 0 |
| Throat foreign sensation | 2 | 0 | 2 | 0 | 0 | 0 |
| Hoarse voice | 3 | 0 | 3 | 0 | 0 | 0 |
| Tiredness Severe fatigue | 3 | 0 | 2 | 0 | 1 | 0 |
| Vomiting | 2 | 0 | 2 | 0 | 0 | 0 |
| Shortness of breath | 2 | 0 | 2 | 0 | 0 | 0 |
| hypofunction | 2 | 0 | 2 | 0 | 0 | 0 |
| Cold sweating | 2 | 0 | 1 | 0 | 1 | 0 |
| Sore eyes | 1 | 0 | 1 | 0 | 0 | 0 |
| Loss or change of sense of taste | 1 | 0 | 1 | 0 | 0 | 0 |
| Abdominal pain | 1 | 0 | 1 | 0 | 0 | 0 |
| Flank pain | 1 | 0 | 1 | 0 | 0 | 0 |
| Lethargy | 1 | 0 | 1 | 0 | 0 | 0 |
| Rhinalgia | 1 | 0 | 1 | 0 | 0 | 0 |
| Lumbago and Low back pain | 1 | 0 | 1 | 0 | 0 | 0 |

SD, standard deviation

### 3.4 Comparison with patient information

Based on a comparison of patient characteristics and SARS-CoV-2 subtypes, we found several correlated features. First, the age distributions of BA.2 patients and BA5 were significantly different. The age distribution of BA.5 patients was younger than that of BA.2 (Figure 3A). Each mutation exhibited a similar pattern of specific mutations (Figure SI5). Second, significant differences were found in some symptoms according to patient age. Patients with fever were younger than those without fever, whereas patients with persistent cough, sneezing, muscle aches, and runny nose tended to be older than those without symptoms (Figure 3B). Finally, differences in the symptoms of each variant were noted. For symptoms of respiratory infection, such as sore throat, persistent cough, sneezing, and runny nose, the ratio of symptoms was higher in BA.2, while systemic symptoms such as fever, headache, muscle aches, and chills were more frequent in BA.5 (Figure 4). However, the relationship between symptom presence and age was observed independently of the virus subtype and age, and the relationship with symptoms was also observed independently of the virus subtype and age.

**Figure 3.**
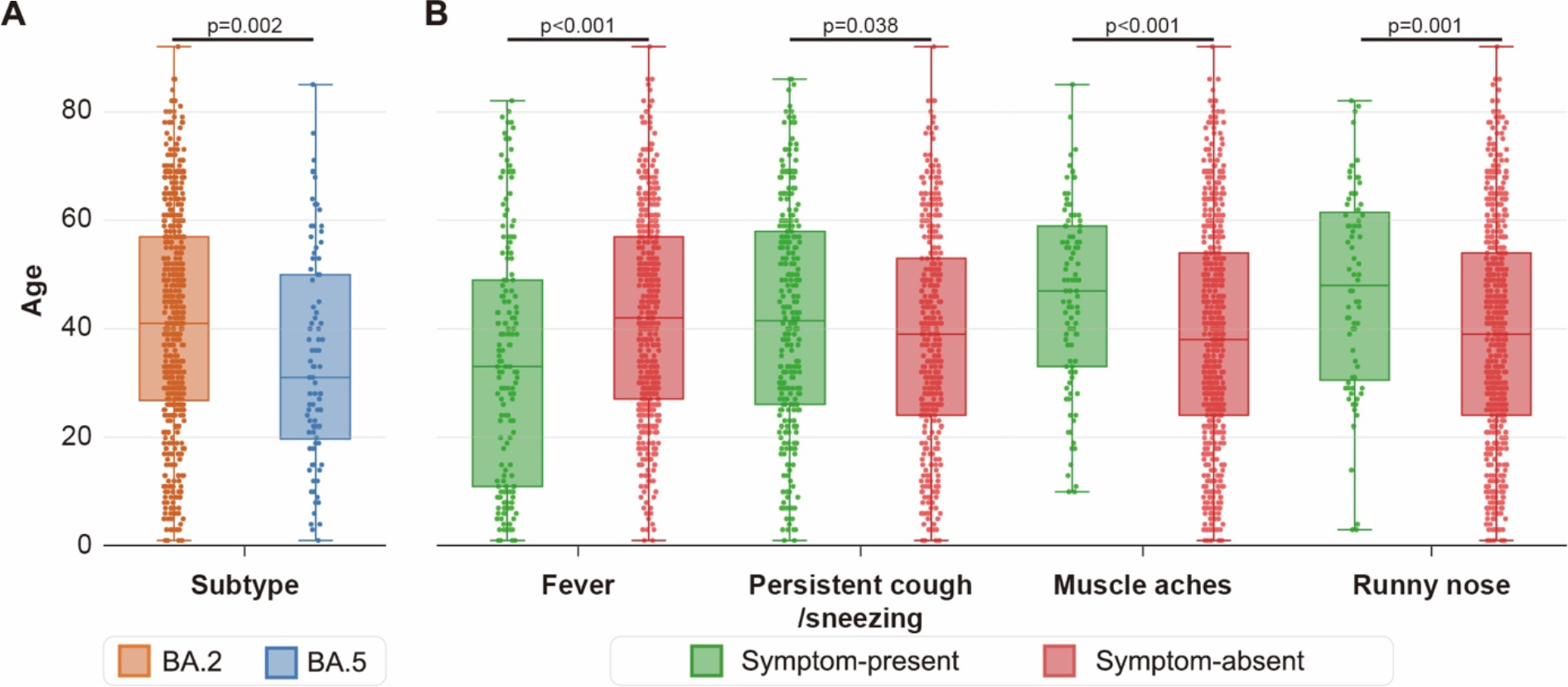
Age distribution by subtype and symptoms. Patient age distribution of BA.2 (red, N=506) and BA.5 (blue, N=87) subtypes is shown in (A). Patient age distribution with or without specific symptoms is also indicated. The number of patients by symptoms is as follows: fever (positive, N=164; negative, N=449), persistent cough & sneezing (positive, N=262; negative, N=351), muscle aches (positive, N=99; negative, N=514), and runny nose (positive, N=73; negative, N=540).

**Figure 4.**
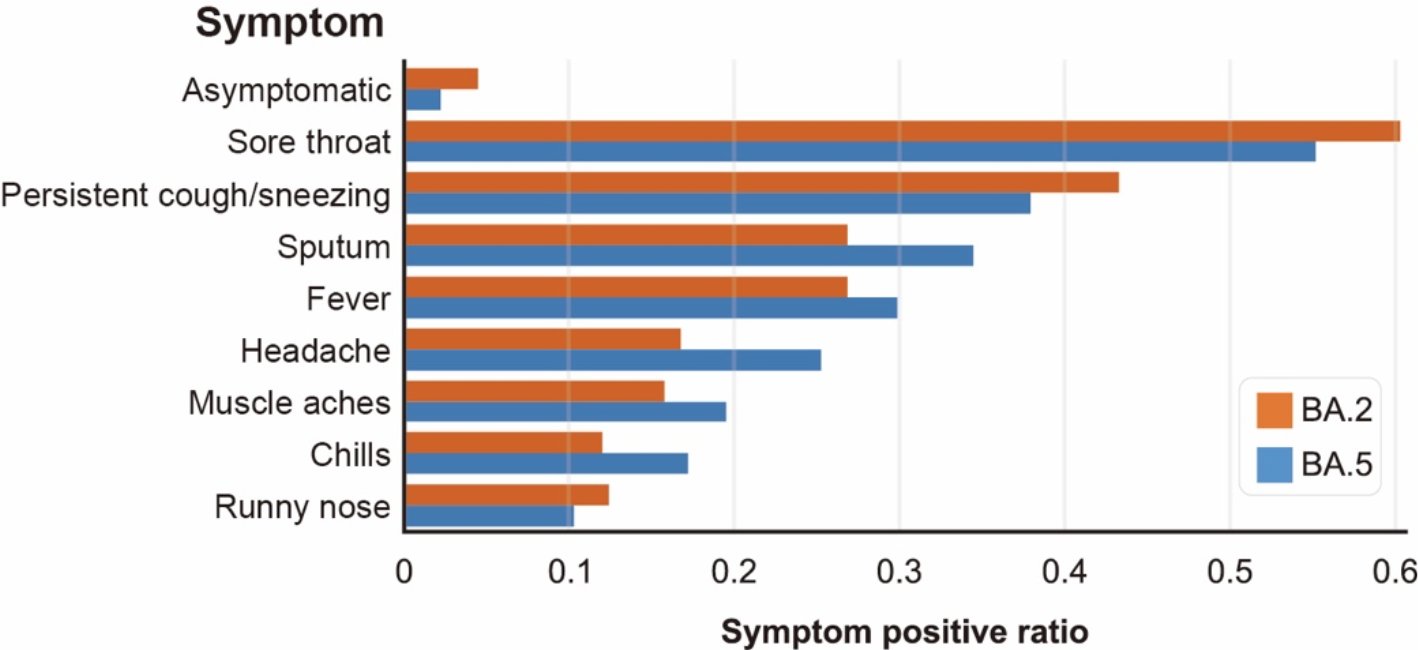
Patient ratio with symptoms by SARS-CoV-2 subtype. For each symptom, patients showed different symptom positive ratio between BA.2 (red, N=506) and BA.5 (blue, N=87). SARS-CoV-2, severe acute respiratory syndrome coronavirus 2

## 4. Discussion

The use of a 2x tiled primer set enabled more accurate detection of SARS-CoV-2 mutations than a single primer set, suggesting that this method may be a promising approach for detecting and monitoring the spread of SARS-CoV-2 and classifying its subtypes. In addition, we applied modified primer composition set similar to one previously implemented to help amplify large deletions in SARS-CoV-2 genome [23, 24]. Furthermore, we highlighted the benefits of employing ONT sequencing for this type of analysis. With longer reads, we obtained complete sequences of the amplified products without assembly and time- or resource-intensive processes. Moreover, ONT sequencing makes the primer set design simpler than other sequencing platforms with shorter read lengths. In addition, while ONT sequencing is known to have a high error rate, amplicon sequencing allowed us to identify the mutations with high accuracy, making it possible to find specific locations and frequencies of SNVs within the SARS-CoV-2 genome [25].

Our findings suggest that the observed differences in symptom profiles and age distributions between BA.2 and BA.5 subtypes may indicate distinct pathogenic mechanisms underlying their transmission and clinical presentation [26, 27]. These findings suggest that different variants may require different diagnostic and treatment approaches to manage symptoms and associated complications. For example, targeted interventions may be necessary to address the specific symptom profiles associated with each variant, which could help improve patient outcomes and reduce the burden on the healthcare system.

In addition to the clinical implications, these findings highlight the importance of ongoing surveillance and monitoring of the SARS-CoV-2 subtypes. As new subtypes emerge and spread, it is critical to understand their pathogenic mechanisms, including differences in symptom profiles and age distributions, to inform public health measures and control the spread of the virus [28]. Further research is necessary to explore the underlying genetic, immune, and environmental factors that contribute to these differences and develop targeted interventions based on this knowledge.

Large-scale studies are necessary to confirm and expand these findings. These studies should include a diverse range of patients with different ages, comorbidities, and symptom profiles to ensure that the results are generalizable and applicable to a wider population. By conducting large-scale studies, researchers can gain a more comprehensive understanding of the pathogenic mechanisms of SARS-CoV-2 and its subtypes.

In conclusion, this study provides important insights into the characteristics of the SARS-CoV-2 subtypes and their impact on patients using long-read ONT sequencing with 2x tiled primer sets. The identification of distinct symptom profiles and age distributions associated with different viral lineages underscores the importance of continued surveillance and monitoring of variants and the need for targeted interventions based on specific symptom profiles. Further research is necessary to confirm and expand on these findings and develop effective strategies to control the spread of SARS-CoV-2 and its subtypes.

## Acknowledgments

We appreciate the patients for their participation and staff at Saint Peter’s Hospital for their support.

## Data Availability

The sequencing data that support the findings of this study are openly available in BioProject at (https://www.ncbi.nlm.nih.gov/bioproject/980725), ID: PRJNA980725.

## References

[1] World Health Organization [Internet]: WHO COVID-19 Dashboard. 2023. Available from: https://covid19.who.int/ x[cited 2023 Apr 20].

[2] Duffy S. Why are RNA virus mutation rates so damn high? PLoS Biol. 2018;16(8):e3000003. doi: 10.1371/journal.pbio.3000003

[3] Sanjuán R, Domingo-Calap P. Mechanisms of viral mutation. Cell Mol Life Sci. 2016;73(23):4433–4448. doi: 10.1007/s00018-016-2299-6

[4] Huang Y, Yang C, Xu XF, et al. Structural and functional properties of SARS-CoV-2 spike protein: potential antivirus drug development for COVID-19. Acta Pharmacol Sin 2020;41(9):1141–1149. 10.1038/s41401-020-0485-4

[5] Shereen MA, Khan S, Kazmi A, et al. COVID-19 infection: Origin, transmission, and characteristics of human coronaviruses. J Adv Res 2020;24:91–98. doi: 10.1016/j.jare.2020.03.005

[6] Candido KL, Eich CR, de Fariña LO, et al. Spike protein of SARS-CoV-2 variants: a brief review and practical implications. Braz J Microbiol. 2022;53(3):1133–5117. doi:10.1007/s42770-022-00743-z

[7] Mengist HM, Kombe AJ, Mekonnen D, et al. Mutations of SARS-CoV-2 spike protein: Implications on immune evasion and vaccine-induced immunity. Semin Immunol. 2021;55:101533. doi: 10.1016/j.smim.2021.101533

[8] Dai L, Gao GF. Viral targets for vaccines against COVID-19. Nat Rev Immunol. 2021;21(2):73–82. doi: 10.1038/s41577-020-00480-0

[9] World Health Organization [Internet]: Classification of Omicron (B.1.1.529): SARS-CoV-2 variant of concern. 2021. Available from: https://www.who.int/news/item/26-11-2021-classification-of-omicron-(b.1.1.529)-sars-cov-2-variant-of-concern; [cited 2021 Nov 26].

[10] Abbasi H, Tabaraei A, Hosseini SM, et al. Real-time PCR Ct value in SARS-CoV-2 detection: RdRp or N gene? Infection. 2022;50(2):537–540. doi: 10.1007/s15010-021-01674-x

[11] Valadan R, Golchin S, Alizadeh-Navaei R, et al. Differential gene expression analysis of common target genes for the detection of SARS-CoV-2 using real time-PCR. AMB Express. 2022;12(1):112. doi: 10.1186/s13568-022-01454-2

[12] Carpenter RE, Tamrakar V, Chahar Het al. Confirming multiplex RT-qPCR use in COVID-19 with next-generation sequencing: strategies for epidemiological advantage. Glob Health Epidemiol Genom. 2022;2022:2270965. doi: 1155/2022/2270965

[13] John G, Sahajpal NS, Mondal AK, et al. Next-generation sequencing (NGS) in COVID-19: a tool for SARS-CoV-2 diagnosis, monitoring new strains and phylodynamic modeling in molecular epidemiology. Curr Issues Mol Biol. 2021;43(2):845–867. doi: 10.3390/cimb43020061

[14] Freed NE, Vlková M, Faisal MB, et al. Rapid and inexpensive whole-genome sequencing of SARS-CoV-2 using 1200 bp tiled amplicons and Oxford Nanopore Rapid Barcoding. Biol Methods Protoc. 2020;5(1):bpaa014. doi: 10.1093/biomethods/bpaa014

[15] Tyson JR, James P, Stoddart D, et al. Improvements to the ARTIC multiplex PCR method for SARS-CoV-2 genome sequencing using nanopore. bioRxiv 2020. 10.1101/2020.09.04.283077

[16] Cheng C, Fei Z, Xiao P. Methods to improve the accuracy of next-generation sequencing. Front Bioeng Biotechnol. 2023;11:982111. doi: 10.3389/fbioe.2023.982111

[17] Grädel C, Terrazos Miani MA, Barbani MT, et al. Rapid and cost-efficient enterovirus genotyping from clinical samples using flongle flow cells. Genes (Basel). 2019;10(9):659. doi: 10.3390/genes10090659

[18] Horiba K, Torii Y, Aizawa Y, et al. Performance of nanopore and Illumina metagenomic sequencing for pathogen detection and transcriptome analysis in infantile central nervous system infections. Open Forum Infect Dis. 2022;9(1):ofac504. doi: 10.1093/ofid/ofac504

[19] Brandt C, Krautwurst S, Spott R, et al. poreCov-an easy to use, fast, and robust workflow for SARS-CoV-2 genome reconstruction via nanopore sequencing. Front Genet. 2021;12:711437. doi: 10.3389/fgene.2021.711437

[20] Robinson JT, Thorvaldsdóttir H, Wenger AM, et al. Variant review with the integrative genomics viewer. Cancer Res. 2017;77(21):e31–e34. doi: 10.1158/0008-5472.CAN-17-0337

[21] Valieris R, Drummond RD, Defelicibus A, et al. A mixture model for determining SARS-Cov-2 variant composition in pooled samples. Bioinformatics. 2022;38(7):1809–1815. doi: 10.1093/bioinformatics/btac047

[22] Thakur P, Thakur V, Kumar P, et al. Emergence of novel omicron hybrid variants: BA(x), XE, XD, XF more than just alphabets. Int J Surg. 2022;104:106727. 10.1016/j.ijsu.2022.106727

[23] Mentes A, Papp K, Visontai D, Stéger J, Csabai I, Medgyes-Horváth A et al. Identification of mutations in SARS-CoV-2 PCR primer regions. Sci Rep 2022;12:18651. doi: 10.1038/s41598-022-21953-3

[24] Mazur-Panasiuk N, Rabalski L, Gromowski T, et al. Expansion of a SARS-CoV-2 Delta variant with an 872 nt deletion encompassing ORF7a, ORF7b and ORF8, Poland, July to August 2021. Euro Surveill. 2021;26(39):2100902. doi: 10.2807%2F1560-7917.ES.2021.26.39.2100902

[25] Player R, Verratti K, Staab A, et al. Comparison of the performance of an amplicon sequencing assay based on Oxford Nanopore technology to real-time PCR assays for detecting bacterial biodefense pathogens. BMC Genomics. 2020;21(1):166. doi: 10.1186/s12864-020-6557-5

[26] Goller KV, Moritz J, Ziemann J, et al. Differences in clinical presentations of Omicron infections with the lineages BA.2 and BA.5 in Mecklenburg-Western Pomerania, Germany, between April and July 2022. Viruses. 2022;14(9):2033. doi: 10.3390/v14092033

[27] Nakakubo S, Kishida N, Okuda K, et al. Associations of COVID-19 symptoms with Omicron subvariants BA.2 and BA.5, host status, and clinical outcomes: a registry-based observational study in Sapporo, Japan. medRxiv. 2023. doi: 10.1101/2023.02.02.23285393

[28] Poovorawan Y, Pyungporn S, Prachayangprecha S, et al. Global alert to avian influenza virus infection: from H5N1 to H7N9. Pathog Glob Health. 2013;107(5):217–223. doi: 10.1179/2047773213Y.0000000103

