## Supplementary Material for "Identification and Analysis of SARS-CoV-2 Mutation and Subtype using 2x tiled Primer Set with Oxford Nanopore Technologies Sequencing for Enhanced Variant Detection and Surveillance in Seoul, Korea"

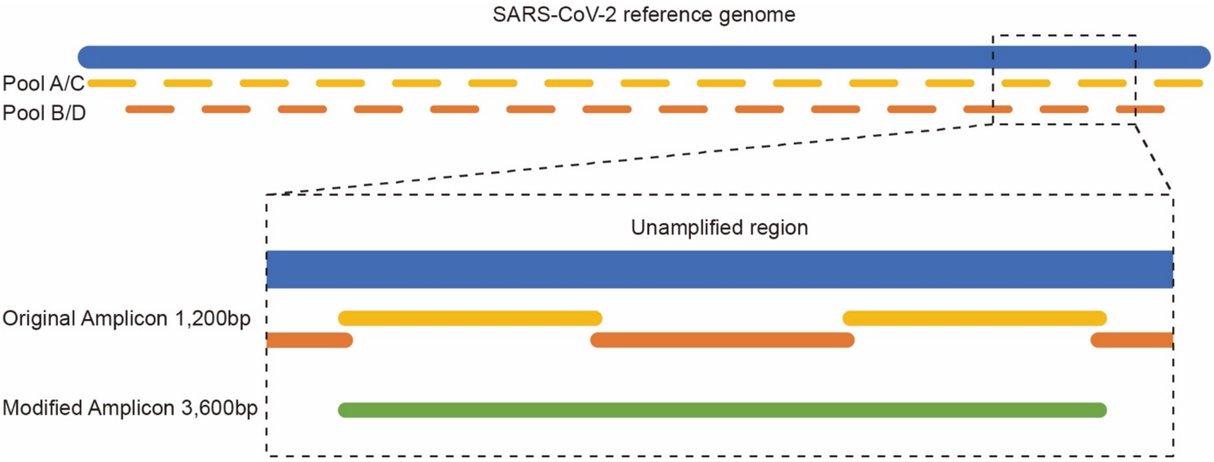


**Supplementary Figure 1. Modified amplicon design for covering large deletions**

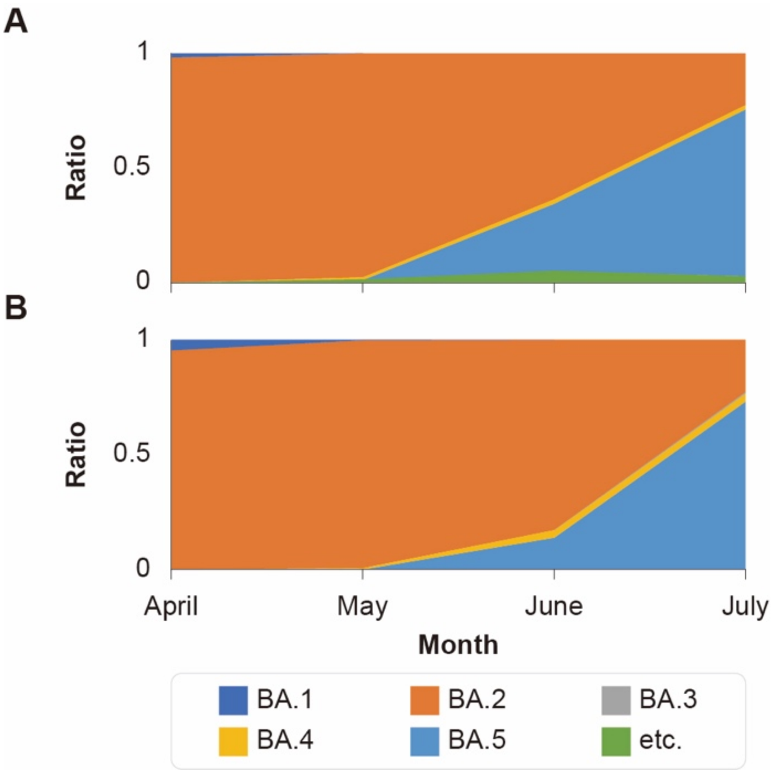


**Supplementary Figure 2. Monthly distribution of SARS-CoV-2 by subtypes in South Korea (April ~ July, 2022)**

(A) Distribution of current Saint Peter's Hospital patients. (B) Distribution from GISAID database. SARS-CoV-2, severe acute respiratory syndrome coronavirus 2

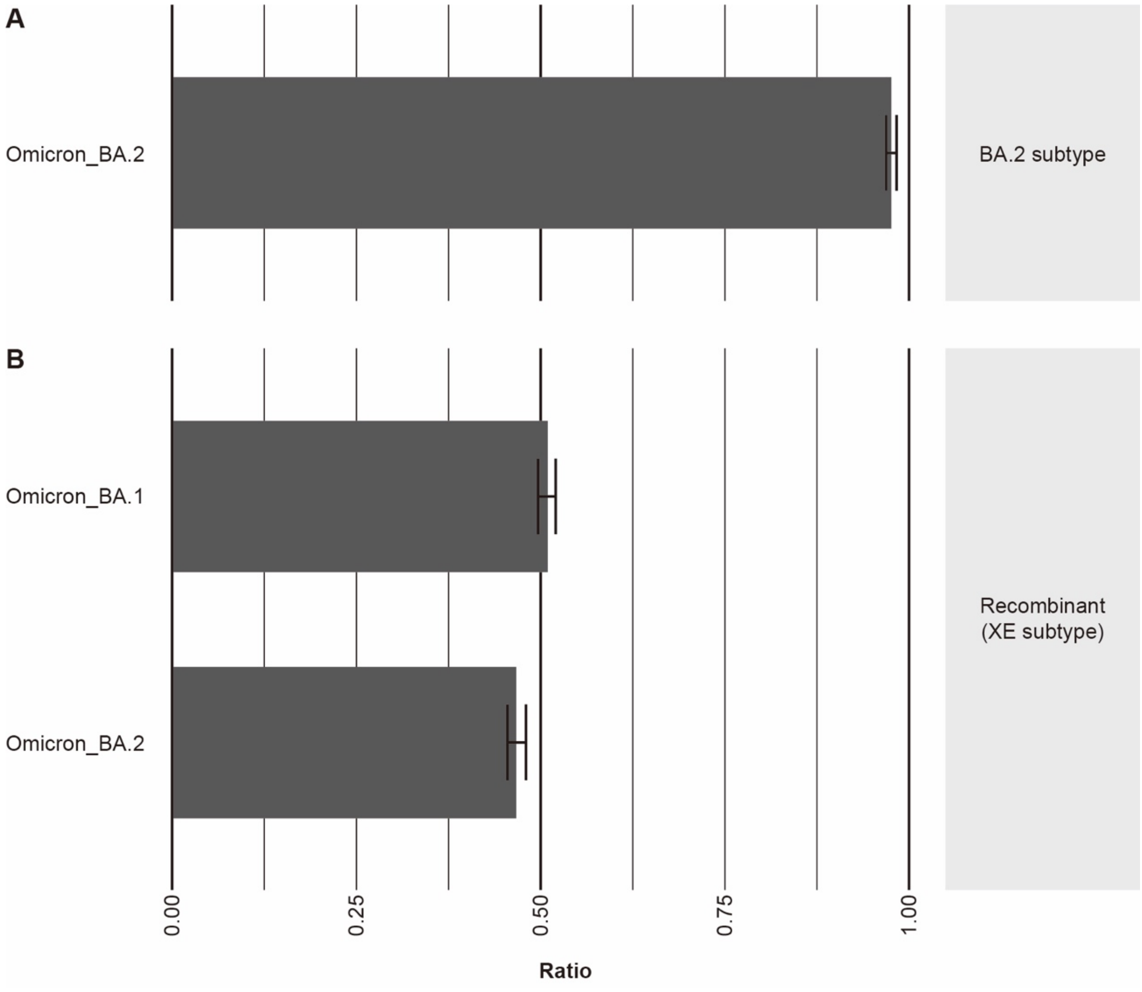


**Supplementary Figure 3. Identification of recombinant ratio by Lineage deComposition for SARS-CoV-2 analysis**

BA.2 subtype samples match with only one subtype (A), and recombinant sample shows mixed ratio over two subtypes (B). SARS-CoV-2, severe acute respiratory syndrome coronavirus 2

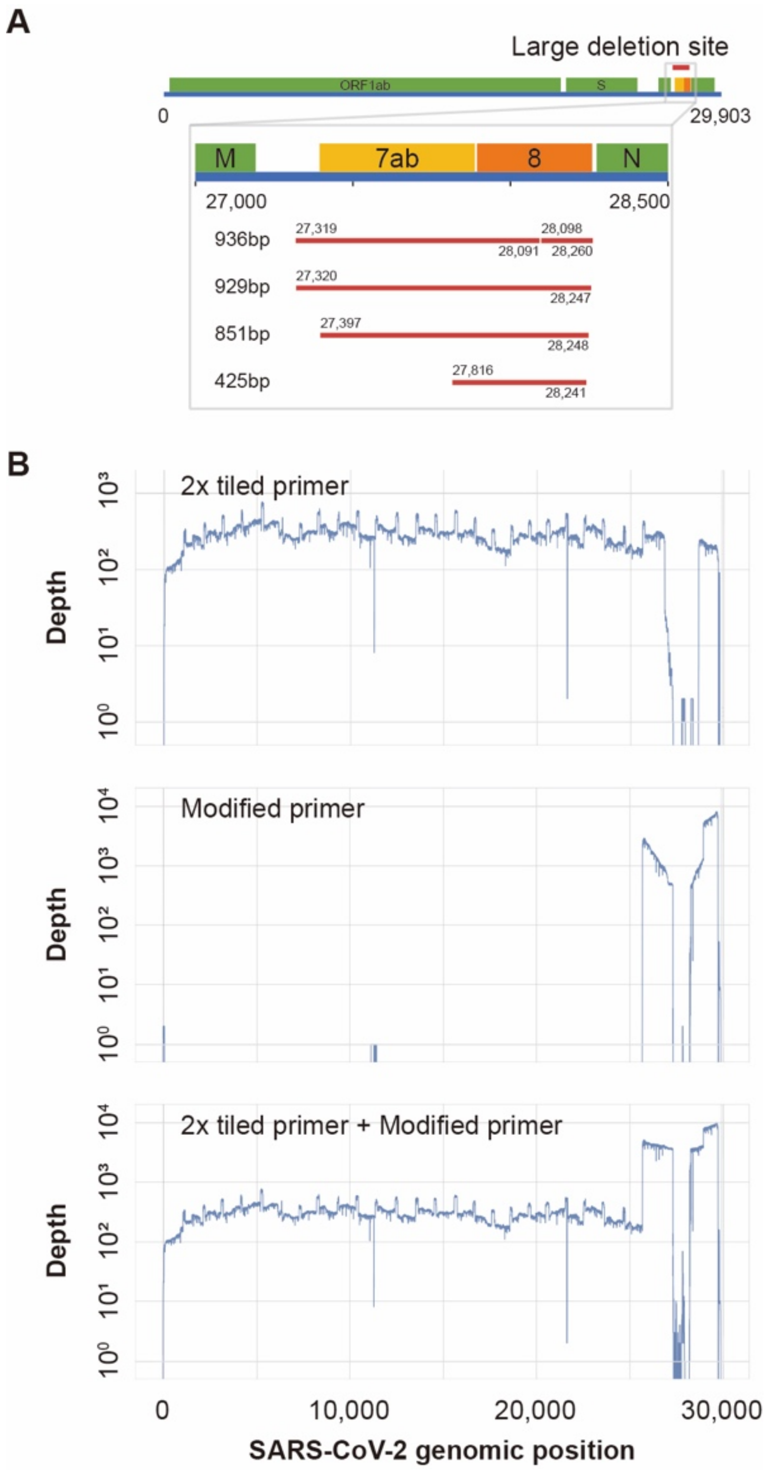


**Supplementary Figure 4. Large deletion of rare SARS-CoV-2 subtype and coverage differences among primer designs**

Large deletions were covered with modified primers and found around ORF7a, ORF7b, and ORF8 (A). Covered region differed by primer design and concatenation of modified primer sequencing data, and 2x tiled primer sequencing data can cover samples with large deletion (B).

SARS-CoV-2, severe acute respiratory syndrome coronavirus 2

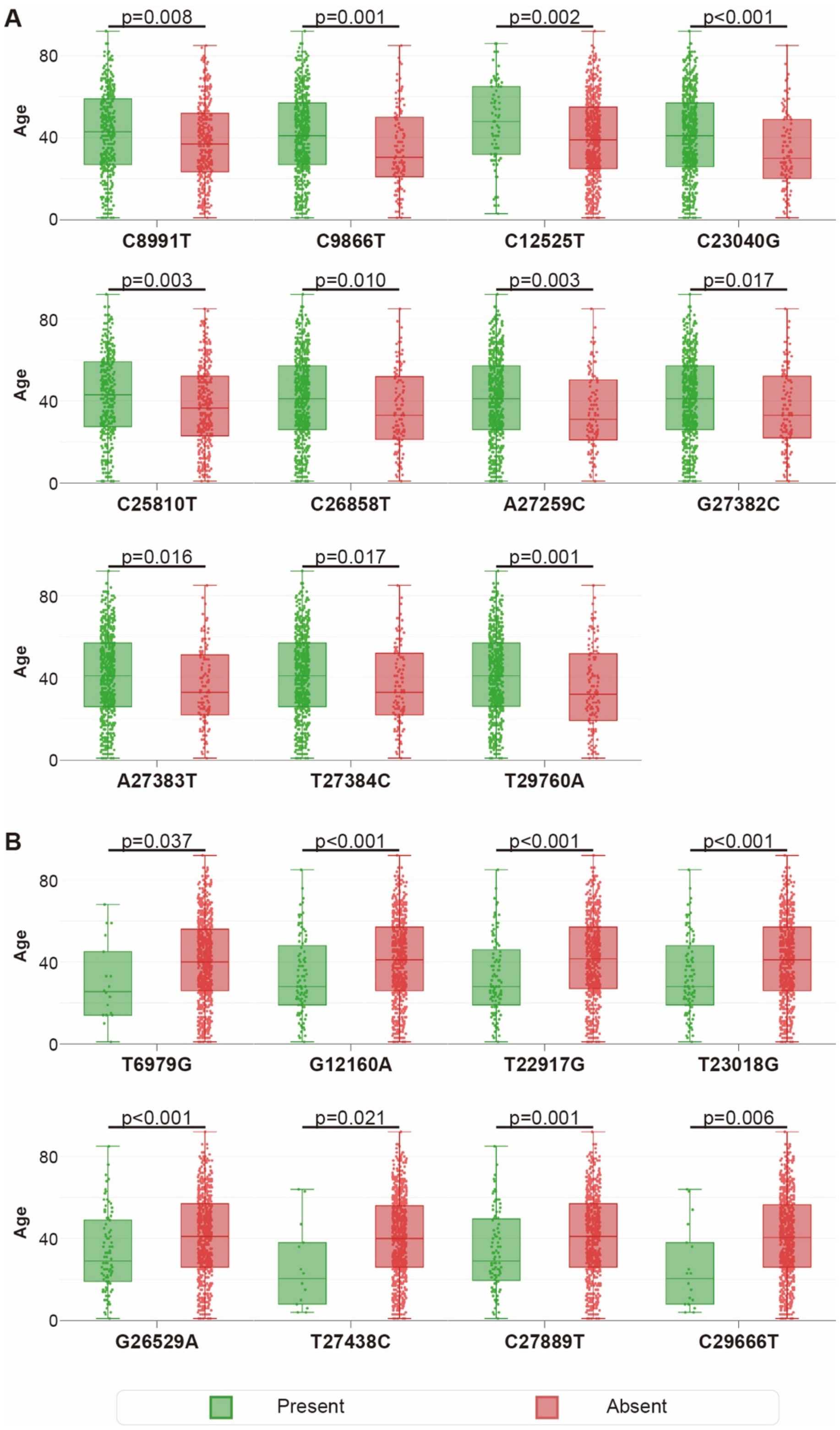


**Supplementary Figure 5. Patient Age distribution by SNVs.**

Age distribution of each subtype-specific SNV also follows subtype age population pattern of BA.2 (A) and BA.5 (B).

SNV, single-nucleotide variations
